# scRGCL: Neighbor-Aware Graph Contrastive Learning for Robust Single-Cell Clustering

**DOI:** 10.64898/2026.03.16.712039

**Authors:** Junming Fan, Fei Liu, Xin Lai

**Affiliations:** School of Software Engineering, South China University of Technology, 510006, Guangzhou, China; Faculty of Medicine and Health Technology,Tampere University, 33520, Tampere, Finland

**Keywords:** single-cell RNA-seq, cell clustering, contrastive learning, systems biology

## Abstract

Accurate cell type identification is a fundamental step in single-cell RNA sequencing (scRNA-seq) data analysis, providing critical insights into cellular heterogeneity at high resolution. However, the high dimensionality, zero-inflated, and long-tailed distribution of scRNA-seq data pose significant computational challenges for conventional clustering approaches. Although recent deep learning-based methods utilize contrastive learning to joint-learn representations and clustering assignments, they often overlook cluster-level information, leading to suboptimal feature extraction for downstream tasks. To address these limitations, we propose scRGCL, a single-cell clustering method that learns a regularized representation guided by contrastive learning. Specifically, scRGCL captures the cell-type-associated expression structure by clustering similar cells together while ensuring consistency. For each sample, the model performs negative sampling by selecting cells from distinct clusters, thereby ensuring semantic dissimilarity between the target cell and its negative pairs. Moreover, scRGCL introduces a neighbor-aware re-weighting strategy that increases the contribution of samples from clusters closely related to the target. This mechanism prevents cells from the same category from being mistakenly pushed apart, effectively preserving intra-cluster compactness. Extensive experiments on fourteen public datasets demonstrate that scRGCL consistently outperforms state-of-the-art methods, as evidenced by significant improvements in normalized mutual information (NMI) and adjusted rand index (ARI). Moreover, ablation studies confirm that the integration of cluster-aware negative sampling and the neighbor-aware re-weighting module is essential for achieving high-fidelity clustering. By harmonizing cell-level contrast with cluster-level guidance, scRGCL provides a robust and scalable framework that advances the precision of automated cell-type discovery in increasingly complex single-cell landscapes.

**Key Messages:**

- scRGCL uses contrastive learning on a regularized representation for single-cell clustering.
- scRGCL outperforms four state-of-the-art methods on 15 datasets.
- scRGCL’s cluster-aware negative sampling and the neighbor-aware re-weighting modules are essential for high-fidelity single cell clustering.

## Introduction

Single-cell RNA sequencing (scRNA-seq) technology overcomes the limitations of traditional bulk sequencing, where the “averaging” effects cellular heterogeneity, by characterizing gene expression at single-cell resolution. Within the scRNA-seq analytical workflow, a critical task involves partitioning cells with similar transcriptomic features into the same cluster [1]. These clustering results not only facilitate the identification of cell subpopulations but also provide foundational support for subsequent downstream analyses, such as elucidating regulatory networks and key signaling pathways [2]. However, constrained by mRNA capture efficiency and the rapid expansion of data scale resulting from reduced sequencing costs, the field continues to face challenges, including high-dimensionality, high sparsity, and noise. Consequently, the development of high-precision clustering models that can effectively mitigate noise while maintaining computational efficiency is of profound significance for deepening the exploration of single-cell transcriptomics underlying cellular heterogeneity and biological processes [3].

scRNA-seq clustering methods can be broadly categorized into traditional machine learning approaches and deep learning–based methods. Traditional approaches typically rely on manual feature engineering and include distance-based partitioning techniques, probabilistic models, and graph-theoretic methods. Early studies such as RaceID [4] employed Pearson correlation coefficients in combination with K-means clustering to identify rare cell populations, while SINCERA [5] established an analysis pipeline centered on hierarchical clustering. To mitigate the challenges posed by high-dimensional data, SC3 [6] applied principal component analysis for feature extraction prior to clustering and incorporated consensus clustering to improve result stability. Graph-based methods, including SNN-Cliq [7] and Sc-GPE [8], further characterized cellular population structures by constructing shared nearest neighbor graphs and applying quasi-clique searching and the Louvain algorithm, respectively.

Traditional scRNA-seq clustering methods provide interpretability but remain sensitive to technical noise. Their dependence on linear dimensionality reduction often results in information loss, failing to account for the complex nonlinear patterns inherent in gene expression. Moreover, these approaches often incur substantial computational overhead and demonstrate restricted scalability when applied to large-scale datasets [9]. Collectively, these limitations motivate the development of more flexible and scalable modeling frameworks capable of robustly learning informative representations from increasingly large and complex single-cell data.

To address these limitations, deep learning has emerged as a powerful alternative for single-cell clustering analysis. Thanks to their hierarchical network structures and strong nonlinear mapping capabilities, deep learning-based approaches take advantage of adaptive feature extraction and effective representation learning. This makes them perfectly suited to addressing the core limitations of traditional methods when processing complex single-cell data. Deep learning–based approaches have thus become the dominant solution for scRNA-seq clustering. Early studies primarily focused on autoencoder-based models such as DCA [10], scDeepCluster [11], scVI [12] and SCVIS [13]. These models perform dimensionality reduction and denoising by explicitly modelling the characteristics of count data through zero-inflated negative binomial (ZINB) distributions or variational inference frameworks. By learning low-dimensional latent embeddings, these methods enable joint optimization of representation learning and clustering, thereby improving robustness to technical noise.

Despite their effectiveness, autoencoder-based methods largely operate on expression profiles alone and often overlook intrinsic relationships among cells. To incorporate structural information, graph neural networks (GNNs) have been introduced to model cell–cell or cell–gene interaction graphs. Representative methods such as GraphSCC [14], graph-sc [15], and scGAC [16] use graph convolutional network (GCN) to capture local topological dependencies. Attention-based variants, including AttentionAE-sc [17], scGNN [18], and scGANSL [19] further incorporate graph attention networks (GATs) to adaptively reweight neighborhood features and better characterize local interactions.

In parallel, contrastive learning has emerged as a complementary paradigm for learning robust representations from noisy single-cell data. Methods such as contrastive-sc [20], scNAME [21], and scGPCL [22] create positive and negative sample pairs using data augmentation or neighborhood-based strategies. This encourages invariance to perturbations and enhances clustering robustness. Nevertheless, challenges remain in designing biologically meaningful augmentations and effectively integrating global dependency modeling with contrastive objectives [23]. Consequently, modern deep clustering frameworks are increasingly oriented toward unifying graph-based modeling, global attention mechanisms, and contrastive learning [24], with the goal of capturing higher-order cellular relationships while maintaining scalability and robustness in large-scale, noisy scRNA-seq datasets.

This study’s objective is to develop a robust framework for single-cell clustering that can model local and global cellular relationships simultaneously, while mitigating the effects of technical noise and heterogeneous cell population densities. To achieve this, we propose scRGCL, a computational framework for the joint optimization of dimensionality reduction and clustering in scRNA-seq data analysis. To overcome the limitation of GAT/GCN capturing only local relationships, we integrate GAT with graph transformer, enabling the realization of “synergistic modeling of local-global structures” by learning a regularized graph. This architecture concurrently captures local micro-topologies and global macro-perspectives, establishing a robust and biologically interpretable computational paradigm for deep mining of scRNA-seq data. To overcome limitations in discriminative power under high technical noise, we introduce contrastive learning leveraging graph edge features. To address heterogeneous density distributions across cell populations, we implement unsupervised clustering using K-means clustering. Together, these integrations (Figure 1) allow scRGCL to achieve a more holistic representation of cellular topology, learn noise-invariant embeddings, and identify rare cell types without pre-defining the number of clusters.

**Figure 1.**
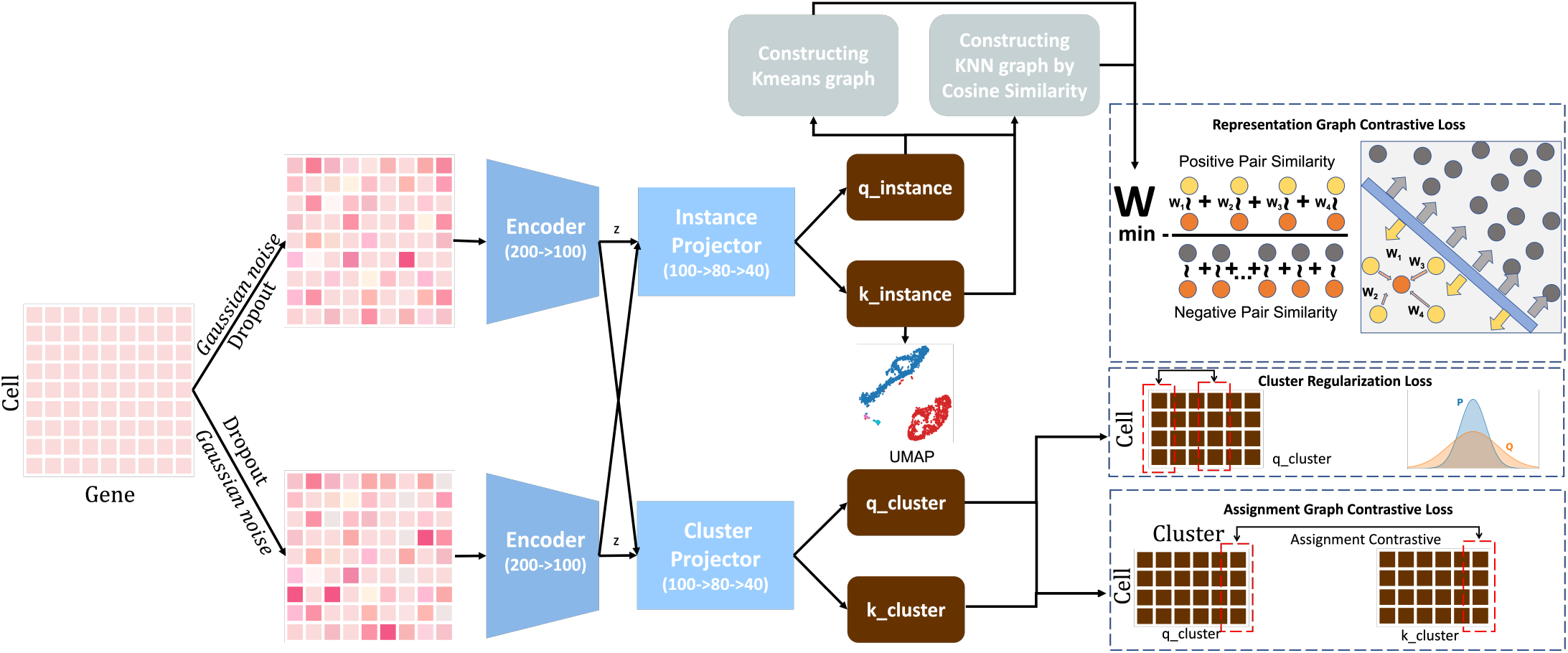
Overall framework of scRGCL. Given a raw gene expression matrix, cells are embedded and used to construct a cell–cell graph based on neighborhood relationships. Two augmented graph views are created via perturbations such as feature masking and Gaussian noise. Each view is encoded by encoder and projector, yielding four latent embeddings. Based on the latent embeddings generated by the Instance Projector, two complementary graphs are constructed: a K-means graph that captures the global clustering structure and a KNN graph—built using cosine similarity—that encodes intricate local neighborhood relationships. The framework optimizes three joint objectives: (a) **Representation Graph Contrastive Loss**: leverages re-weighted noise contrastive estimation loss to enforce augmented features of connected cells in the graph to be similar and disconnected ones to be distinct; (b) **Cluster Regularization Loss**: is defined as the difference between the logarithm of cluster number and the entropy of cluster size distribution, (c) **Assignment Graph Contrastive Loss**: imposes cluster-level consistency on assignment probabilities by aligning the cluster assignment distribution of augmented cells with that of their random graph neighbors. The learned embeddings are employed for downstream tasks such as clustering.

## Methods

### Problem Definition

Given a single-cell RNA sequencing dataset consisting of *N* cells, we denote the gene expression matrix as *X* = {*x*_1_, *x*_2_, …, *x*_*N*_ } ∈ ℝ^*N ×G*^, where *G* represents the number of genes. The objective of our method is to partition these *N* cells into *K* distinct clusters based on their semantic similarities, without the guidance of ground-truth labels. Unlike image data, scRNA-seq data is characterized by high dimensionality, sparsity, and technical noise (e.g., dropout events). Therefore, we adapt the Graph Contrastive Clustering (GCC) framework to effectively capture both local neighborhood information and global cluster structures in the single-cell domain.

### Data Preprocessing

To ensure the quality of the input data, we perform standard preprocessing on the raw count matrix *X*. To ensure data quality and computational stability, cells and genes with zero total counts were filtered out to eliminate uninformative observations. The library size of each cell is normalized to 10,000 counts, followed by a logarithmic transformation *x*^*′*^ = log(*x* + 1). Finally, we select the top 2000 highly variable genes (HVGs) to form the input feature vectors for the network.

### Data Augmentation

Contrastive learning relies on constructing positive pairs via data augmentation. Since geometric transformations (e.g., cropping, flipping) used in imaging processing are inapplicable to gene expression vectors, we employ biologically relevant augmentation strategies

- **Bernoulli Masking (Dropout Simulation):** To mimic the technical dropout noise in scRNA-seq, we randomly mask a subset of non-zero gene expression values to zero with a probability *p*_*mask*_ = 0.2.
- **Gaussian Noise Injection:** We inject random Gaussian noise *ϵ* ∼ 𝒩 (0, *σ*^2^) into the expression profile to simulate data variability. For a given cell *x*_*i*_, these augmentations generate a transformed view 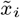, enforcing the model to learn representations invariant to technical noise.

### Graph Construction

The core of scRGCL utilizes topological structures to guide contrastive learning. We construct a graph *G* = (*V, E*) where vertices *V* represent cells and the edge set *E* encodes the neighboring relationships between cells. Specifically, *E* is constructed by first applying k-means clustering to group cells into preliminary clusters, followed by building a k-nearest neighbor (KNN) graph based on cosine similarity within the learned embedding space to capture local cell-to-cell similarities. To mitigate the bias caused by fluctuating features during training, we utilize a moving average of the representation features *z*.

The adjacency matrix *A* is constructed using the KNN algorithm with *k* values ranging from 4 to 10 based on dataset characteristics:

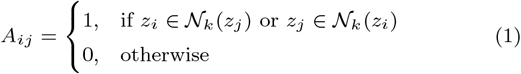

where 𝒩_*k*_(*z*_*i*_) denotes the set of *k*-nearest neighbors of cell *i*. The normalized graph Laplacian is subsequently computed as *L* = *I − D*^*−*1/2^ *AD*^*−*1/2^, where *D* is the degree matrix with *D*_*ii*_ = Σ_*j*_ *A*_*ij*_ .

### Graph Contrastive Clustering Framework

Our framework employs a dual-head architecture sharing a backbone encoder *f*_*θ*_ (e.g., a Multi-Layer Perceptron or Graph Attention Network). The encoder maps the input *x*_*i*_ to a normalized feature representation *z*_*i*_ ∈ ℝ^*d*^ and a cluster assignment probability vector *p*_*i*_ ∈ ℝ^*K*^ . The training objective integrates two graph-based contrastive modules.

### Representation Graph Contrastive (RGC)

The RGC module aims to learn discriminative features by pulling connected neighbors together in the representation space. Unlike standard instance discrimination, which treats all other samples as negatives, RGC considers graph neighbors as positive samples. The loss function is defined as:

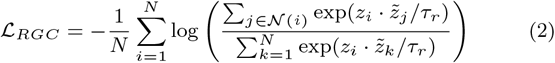

where 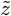 represents the feature of the augmented view, 𝒩(*i*) is the neighbor set of sample *i* defined by the graph, and *τ*_*r*_ is the temperature parameter. Minimizing this loss reduces the intra-community variance of features.

### Assignment Graph Contrastive (AGC)

To achieve cluster-level consistency, the AGC module enforces that a cell and its neighbors should share similar cluster assignment distributions. Let *q*_*c*_ denote the column vector of the assignment matrix corresponding to cluster *c* (i.e., the probability distribution of all samples over cluster *c*). The AGC loss maximizes the agreement between the cluster assignments of the original graph and the augmented view:

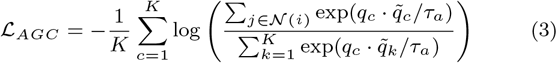

where *τ*_*a*_ is the assignment temperature. This module ensures that the learned clusters are compact and consistent with the underlying manifold structure.

### Overall Objective

To prevent the trivial solution where all samples are assigned to a single cluster, we incorporate a cluster regularization term ℒ_*CR*_ based on the entropy of the assignment probabilities:

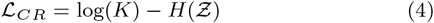

where *H* denotes entropy,

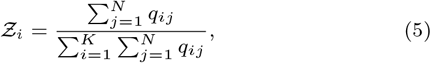

and **q** = [*q*_1_, …, *q*_*K*_ ]_*N×K*_ is the assignment probability matrix with *q*_*ij*_ denoting the probability that sample *j* is assigned to cluster *i*. The final objective function is a weighted sum of the three components:

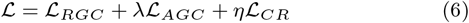

where *λ* and *η* are hyperparameters balancing the contribution of assignment consistency and regularization.

### Evaluation Metrics

To assess clustering performance, we utilized two widely adopted external validation metrics: **Adjusted Rand Index (ARI)** and **Normalized Mutual Information (NMI). ARI** evaluates pairwise consistency between partitions while correcting for chance agreement. It ranges from *−*1 to 1, where 1 represents perfect alignment and 0 corresponds to random labeling. **NMI** quantifies the shared information between predicted clusters *C* and ground-truth labels *Y*, normalized by their respective entropies. It ranges from 0 to 1, with higher values indicating stronger semantic agreement. All metrics were computed using the `cikit-learn` library.

## Results

scRGCL integrates graph contrastive learning with cluster-aware sampling to achieve robust scRNA-seq clustering. Starting from a raw expression matrix, scRGCL generates augmented views to mitigate technical noise and extracts low-dimensional embeddings via a residual-attention autoencoder. By incorporating graph-aware contrastive learning, scRGCL preserves local neighborhood structures while ensuring global consistency across clusters. Finally, the learned embeddings are used for unsupervised clustering, enabling accurate identification of cell populations.

To comprehensively evaluate the performance of *scRGCL*, we conducted extensive experiments on a benchmark comprising 15 publicly available scRNA-seq datasets (**Supplementary Table S1**). These datasets encompass diverse tissue types, sequencing technologies, and species, enabling thorough validation of our method’s generalization capability under varying conditions. The 15 datasets cover multiple tissues (e.g., bladder, muscle, spleen, diaphragm, trachea, brain, pancreas, breast), species (mouse and human), and protocols (10x Genomics, Smart-seq2, and inDrop), including multi-batch pancreas cohorts (Baron1–4). Standard single-cell data preprocessing was performed, including quality control, library size normalization, log-transformation, and selection of the top 2,000 highly variable genes using Seurat(see Methods).

### Overall Performance Comparison

We compared our method with four representative baselines: scCCL [25] (contrastive learning with momentum encoder and dual-space instance/cluster-level learning), scLEGA [26] (dual-branch DAE-GAE with attention fusion and ZINB loss), scSAMAC [27] (VAE with saliency-adjusted masking and Wasserstein clustering), and AttentionAE-sc [17] (attention-based autoencoder with DEC). Among these, scCCL is most closely related to our work as both leverage contrastive learning; the others are autoencoder-based methods included to examine whether source-derived negatives benefit clustering within that paradigm. Detailed descriptions of the baseline methods are provided in **Supplementary Materials**. *scRGCL* demonstrated superior clustering performance across all 15 datasets, achieving an average ARI of 89.35% and average NMI of 83.41%, significantly outperforming all baseline methods (Table 1). In ARI metrics, *scRGCL* improved by 8.34% over the second-best method scCCL (81.01%), and by 16.96%, 17.09%, and 16.09% over scLEGA (72.39%), scSAMAC (72.26%), and scAttentionAE (73.26%), respectively. In NMI metrics, improvements were similarly substantial, with *scRGCL* achieving 83.41% compared to scCCL’s 79.42%, representing a 3.99% improvement. These results show that our regularized graph contrastive learning framework effectively integrates local neighborhood information with global clustering structures to learn more discriminative cellular representations.

We further analyzed the scaling behavior of methods across datasets of varying sizes. *scRGCL* demonstrated stable high performance on small-scale (e.g., Pollen containing 301 cells), medium-scale (e.g., Quake_Smart-seq2_Diaphragm containing 870 cells), and large-scale (e.g., Quake_10x_Spleen containing 9,552 cells) datasets. This scaling property is significant for practical applications involving datasets of different magnitudes. In contrast, the baseline methods showed performance variation on all datasets. Specifically, *scRGCL* achieved an average ARI of 89.35% with a standard deviation of 9.21%, while scCCL achieved an average ARI of 81.01% with a standard deviation of 20.45%. This comparison indicates that *scRGCL* not only achieves higher average performance but also exhibits lower performance variance across different data distributions, confirming its stability and robustness.

**Table 1.** Clustering performance comparison on benchmark datasets. Performance is measured by ARI (%). Best results are shown in bold. Datasets with missing values (“–”) indicate that the corresponding method failed to converge due to gradient explosion. For compactness, we abbreviate `Quake_10x` as `Q10x` and `Quake_Smart-seq2` as seq2.

| Dataset | ARI (%) |  |  |  |  | NMI (%) |  |  |  |  |
| --- | --- | --- | --- | --- | --- | --- | --- | --- | --- | --- |
|  | scRGCL | scCCL | scLEGA | scSAMAC | scAttnAE | scRGCL | scCCL | scLEGA | scSAMAC | scAttnAE |
| Baron1 | 87.98 | **92.87** | 92.46 | 73.52 | 68.95 | 79.47 | 84.77 | **89.21** | 78.84 | 79.52 |
| Baron2 | **90.58** | 89.69 | 12.14 | 89.84 | 85.57 | **85.01** | 87.11 | 35.84 | 82.69 | 80.22 |
| Baron3 | 88.76 | 86.88 | 85.18 | **91.20** | 88.80 | 85.04 | 84.11 | 86.06 | **86.35** | 86.35 |
| Baron4 | **86.79** | 81.40 | 87.20 | 85.44 | 85.30 | 80.25 | 82.54 | **86.82** | 82.45 | 86.76 |
| Chung | **62.85** | 22.05 | 41.17 | 14.72 | 17.71 | **45.30** | 15.50 | 48.74 | 16.42 | 37.19 |
| Klein | **88.40** | 78.38 | 75.58 | 86.54 | 73.40 | **84.69** | 78.04 | 76.18 | 89.93 | 74.51 |
| Muraro | 90.47 | 91.94 | **93.51** | 83.75 | 92.35 | 84.21 | 87.05 | **89.71** | 80.51 | 88.31 |
| Pollen | **93.79** | 75.30 | 55.66 | 25.36 | 53.38 | **91.81** | 83.89 | 77.24 | 36.35 | 76.31 |
| Q10x_Bladder | **98.81** | 75.52 | 50.95 | 61.98 | 76.76 | **95.79** | 80.71 | 69.14 | 73.79 | 82.02 |
| Q10x_Limb_Muscle | 99.17 | **99.43** | 97.63 | 86.27 | 97.67 | 98.31 | **98.85** | 95.03 | 91.41 | 94.82 |
| Q10x_Spleen | 89.77 | **90.02** | 82.51 | – | 73.98 | 78.80 | **79.33** | 77.17 | – | 7116 |
| Romanov | **78.30** | 75.54 | 58.50 | 59.99 | 77.58 | **70.48** | 68.60 | 65.72 | 68.08 | 71.94 |
| seq2_Diaphragm | 97.42 | 98.29 | 97.04 | 44.27 | 91.45 | 94.91 | **96.11** | 93.55 | 55.59 | 89.18 |
| seq2_Limb_Muscle | 93.11 | 97.82 | 95.32 | 72.19 | 79.16 | 86.11 | **95.74** | 90.84 | 61.24 | 84.18 |
| seq2_Trachea | **94.01** | 60.06 | 61.02 | 71.89 | 63.42 | **90.97** | 68.93 | 73.25 | 52.37 | 72.24 |
| Average | **89.35** | 81.01 | 72.39 | 67.64 | 75.03 | **83.41** | 79.42 | 76.97 | 68.29 | 78.31 |
| Standard Deviation | 9.04 | 19.62 | 25.04 | 24.22 | 19.72 | 12.86 | 19.71 | 16.85 | 21.84 | 13.41 |

### Ablation Study

To understand the contribution of each module to *scRGCL*’s performance, we designed systematic ablation experiments evaluating the effects of instance-level contrast, cluster-level contrast, and cluster regularization mechanisms. As a result, the ablation study covered four model variants: the complete model (All), a variant without the representation graph contrastive module (no RGC), a variant without the assignment graph contrastive module (no AGC), and a variant without the cluster regularization module (no CR). The ablation results indicate that the representation graph contrastive module is the core performance driver of *scRGCL* (Table 2). Removing it (no RGC) reduced the average ARI from 89.35% to 65.77%, the largest drop among all variants. This aligns with its role in shaping discriminative embeddings. On some datasets (e.g., Baron1), the degradation was particularly severe, dropping from 87.98% to 45.09%, underscoring the importance of this module for heterogeneous data. Removing the assignment graph contrastive module (no AGC) reduces the average ARI to 80.40% by 8.95%, indicating that this component improves inter-cluster separation and compactness via higher-level consistency constraints. Removing cluster regularization module (no CR) also lowers the average ARI to 77.72% by 11.63%, highlighting its role in mitigating class imbalance by up-weighting samples from rare cell types and preventing them from being marginalized during contrastive learning. Removing these modules has similar impacts on the model’s NMI values.

**Table 2.** Ablation study results on benchmark datasets. Performance is reported as ARI (%) and NMI (%) for the full model (ALL) and variants removing the representation graph contrastive (no RGC), assignment graph contrastive (no AGC), or cluster regularization (no CR) module. Best results are shown in bold. For compactness, we abbreviate `Quake_10x` as `Q10x` and `Quake_Smart-seq2` as seq2.

| Dataset | ARI (%) |  |  |  | NMI (%) |  |  |  |
| --- | --- | --- | --- | --- | --- | --- | --- | --- |
|  | ALL | no RGC | no AGC | no CR | ALL | no RGC | no AGC | no CR |
| Baron1 | **87.98** | 45.09 | 73.42 | 60.97 | **79.47** | 63.06 | 73.49 | 76.82 |
| Baron2 | **90.58** | 54.09 | 72.44 | 73.06 | **85.01** | 71.87 | 74.24 | 70.40 |
| Baron3 | 88.76 | 65.59 | 86.28 | **91.04** | 85.04 | 73.74 | 80.89 | **85.89** |
| Baron4 | **86.79** | 60.29 | 69.49 | 70.56 | **80.25** | 74.25 | 79.71 | 75.16 |
| Chung | **62.85** | 47.89 | 48.67 | 39.20 | **45.30** | 35.09 | 27.52 | 41.59 |
| Klein | 88.40 | 67.42 | 92.21 | **92.85** | 84.69 | 65.85 | 87.91 | **89.18** |
| Muraro | **90.47** | 74.88 | 76.02 | 73.66 | **84.21** | 68.63 | 82.27 | 79.16 |
| Pollen | **93.79** | 33.14 | 76.82 | 83.55 | **91.81** | 40.33 | 78.92 | 83.53 |
| Q10x_Bladder | 98.81 | 98.48 | **99.23** | 76.31 | 95.79 | 94.82 | **96.53** | 81.01 |
| Q10x_Limb_Muscle | **99.17** | 85.61 | 95.59 | 95.07 | **98.31** | 84.15 | 94.93 | 94.38 |
| Q10x_Spleen | **89.77** | 75.58 | 75.03 | 89.54 | **78.80** | 62.59 | 61.97 | 78.27 |
| Romanov | **78.30** | 68.05 | 74.12 | 75.12 | 70.48 | 53.32 | 66.62 | **71.61** |
| seq2_Diaphragm | **97.42** | 72.35 | 94.54 | 97.09 | **94.91** | 64.92 | 89.92 | 94.59 |
| seq2_Limb_Muscle | **93.11** | 82.39 | 86.39 | 87.13 | **86.11** | 78.16 | 82.11 | 82.23 |
| seq2_Trachea | **94.01** | 55.67 | 85.69 | 60.67 | **90.97** | 45.68 | 76.76 | 53.09 |
| Average | **89.35** | 65.77 | 80.40 | 77.72 | **83.41** | 65.10 | 76.92 | 77.13 |

### t-SNE Visualization

We used t-SNE plots to qualitatively evaluate cluster quality across diverse datasets (Figure 2). While autoencoder-based baselines often produce hyper-compact, spherical clusters through aggressive latent compression, they tend to sacrifice transitional biological information and manifold topology. In contrast, contrastive learning methods like scRGCL optimize relative distances, allowing clusters to naturally conform to the data’s intrinsic distribution. Notably, compared to the contrastive baseline scCCL, scRGCL demonstrates a superior identification of fine-grained cell types; it effectively resolves distinct subpopulations even within continuous, “bridged” clusters where expression profiles highly overlap. These visual patterns—characterized by high fidelity to ground truth and clear inter-cluster margins—align with our quantitative gains in ARI/NMI, confirming that scRGCL excels at capturing both global structure and delicate biological boundaries.

**Figure 2.**
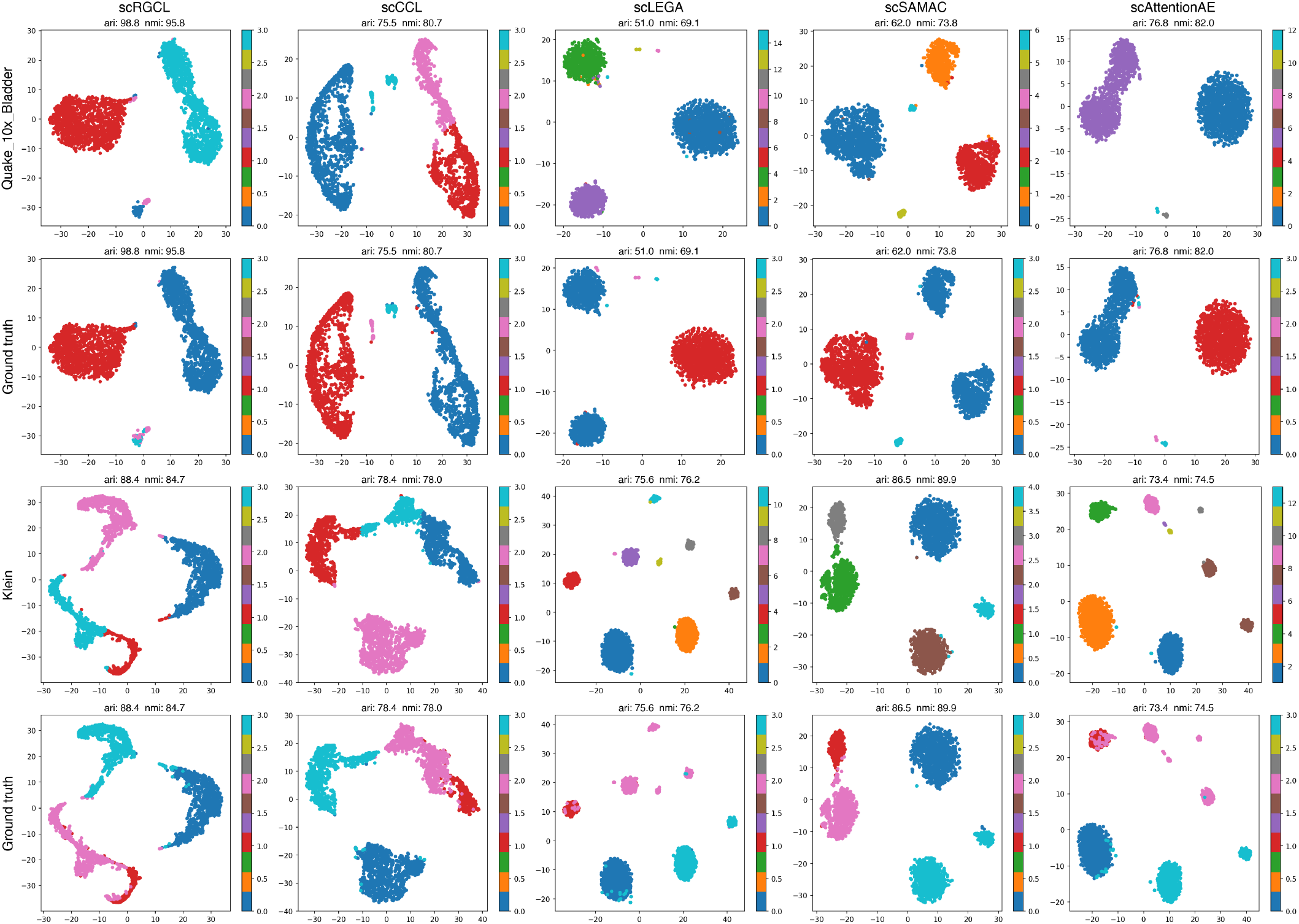
t-SNE visualization of clustering results on representative datasets. Rows correspond to datasets (Quake 10x Bladder, Ground truth (Quake 10x Bladder), Klein, Ground truth (Klein)) and columns correspond to methods (scRGCL, scCCL, scLEGA, scSAMAC, scAttentionAE). Note that since clustering is an unsupervised process, the categorical color assignments for predicted labels are arbitrary and may not strictly correspond to the specific colors used for ground truth labels.

The corresponding visualization results for the remaining datasets are presented in **Supplementary Fig. S1–S3**.

## Conclusion

### Summary

This work proposes scRGCL, a graph contrastive learning framework for robust clustering of scRNA-seq data with high dimensionality, sparsity, and technical noise. scRGCL unifies local and global structure modeling by combining neighborhood-aware graph learning (via representation graph contrastive) with global clustering consistency (via assignment graph contrastive), enabling the model to capture both short-range micro-topologies and long-range cellular dependencies. In addition, the representation graph contrastive and assignment graph contrastive objectives promote noise-invariant embeddings and cluster-consistent assignments, improving stability under dropout and other technical perturbations.

### Discussion

Despite its robust performance, scRGCL has limitations that need future exploration. The current dependency on a pre-specified cluster number may constrain the discovery of novel cell types in exploratory analyses, while the sensitivity of KNN graph construction to severe batch effects can introduce noisy edges that impact regularization. Future work will focus on developing adaptive mechanisms for cluster determination and refining graph construction algorithms to enhance the model’s generalization and stability in the presence of systemic technical biases.

## Data availability

The code and data used in this article are available in a GitHub repository https://github.com/laixn/scRGCL.

## Author contributions statement

JF: methodology; software; data curation; writing–original draft; writing–review and editing. FL: methodology; supervision; writing–review and editing. XL: conceptualization; methodology; software; data curation; investigation; validation; formal analysis; supervision; funding acquisition; visualization; project administration; resources; writing–original draft; writing–review and editing.

### Abbreviations

scRNA-seq: single-cell RNA sequencing
HVGs: Variable Genes
ZINB: zero-Inflated Negative Binomial

